# Structural and biochemical analysis of the IBV nsp15 endoribonuclease reveals the necessity of peripheral site residues for activity

**DOI:** 10.64898/2026.08.28.747898

**Authors:** Ena S. Tully, Robert N. Kirchdoerfer

## Abstract

Infectious bronchitis virus (IBV) is a member of the *Gammacoronavirus* genus responsible for respiratory illness and weakened eggshells in infected chickens, adversely impacting the poultry industry. Escaping innate immune detection during infection is crucial for coronavirus proliferation in the host. The production of double-stranded RNA during coronavirus replication triggers innate immune sensors to create an antiviral state within infected cells. To counter this response, coronaviruses employ nonstructural protein 15 (nsp15) endoribonuclease to degrade double-stranded RNA. Here, we use cryo-electron microscopy and biochemistry to characterize IBV nsp15’s interactions with RNA. While the overall structure and active site of IBV nsp15 strongly resemble previous studies of nsp15 from other coronaviral genera, we note that double-stranded RNA contacts several non-conserved residues peripheral to the enzyme active site. Our data show that these residue positions can have strong impacts on RNA cleavage suggesting unique solutions for RNA engagement across coronavirus species. We also demonstrate a preference for IBV nsp15 to cleave double-stranded RNA over single-stranded RNA and observe nsp15 hexamers with two double-stranded RNAs bound simultaneously. This study reenforces the need to study diverse coronavirus species to identify distinct viral enzyme characteristics.

**GRAPHICAL ABSTRACT:** 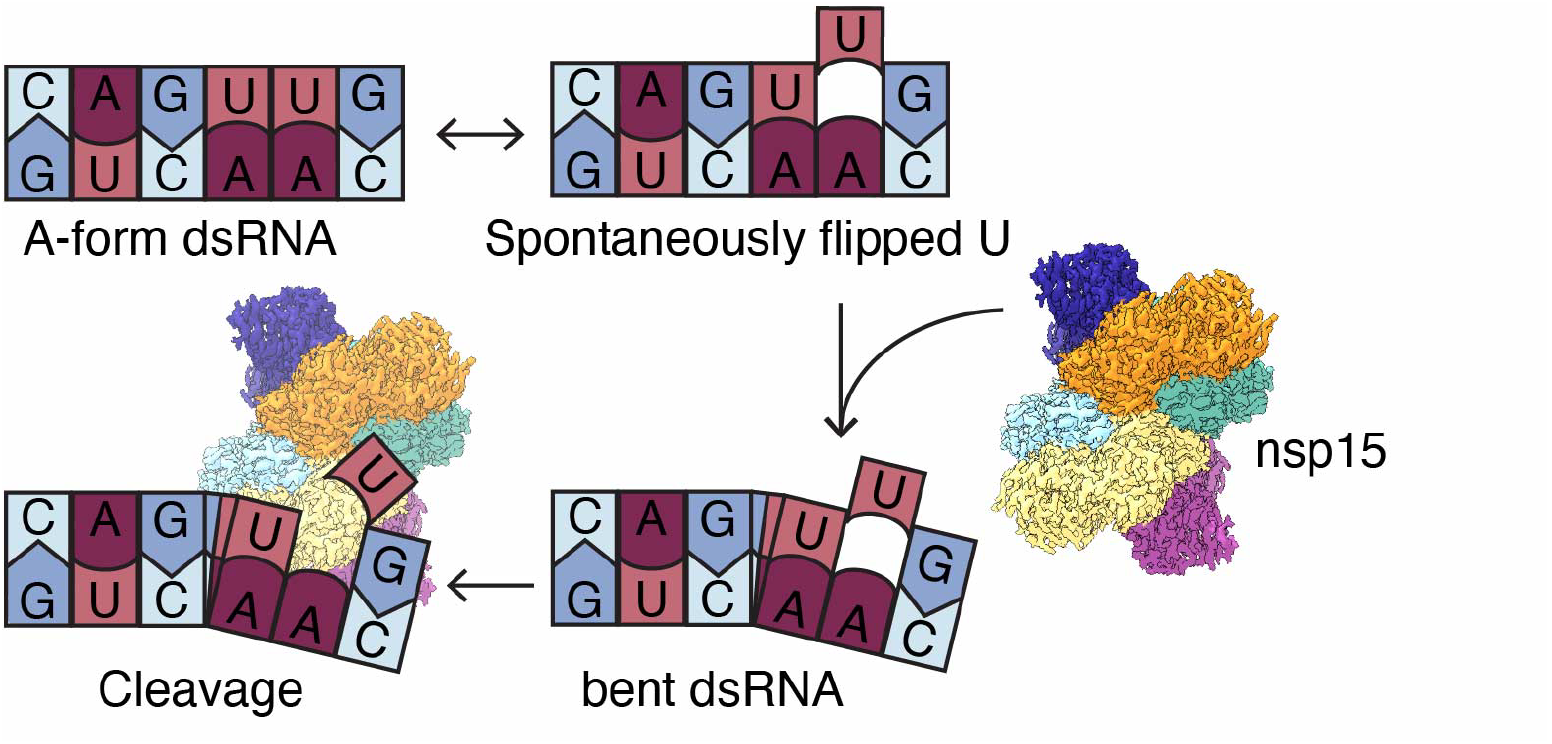

## INTRODUCTION

Coronaviruses are a group of positive-sense single-stranded RNA (ssRNA) viruses within the *Nidovirales* order and the *Coronaviridae* family and are divided into the *Alphacoronavirus, Betacoronavirus, Gammacoronavirus*, and *Deltacoronaviurs* genera. Coronaviruses infect diverse animal hosts. Crossover events from animal hosts to humans have led to outbreaks and pandemics which have had detrimental impacts on human health (1). Starting with SARS-CoV in 2002, followed by MERS-CoV in 2012, and most recently SARS-CoV-2 in 2019, highly pathogenic coronaviruses have emerged approximately every decade, leading to speculation of future viral emergences (2).

While many coronaviruses have had significant impacts on human health, most coronaviruses infect non-human animal hosts. Once such virus, infectious bronchitis virus (IBV) of the *Gammacoronavirus* genus, is a highly transmissible virus that creates a significant burden to the poultry industry, resulting in a yearly economic loss estimated to be $3 billion worldwide and causing a highly pathogenic respiratory disease in chickens (3). Economic costs and losses are associated with vaccine regimen implementation, treatment of infected animals, culling infected chickens, and losses in egg production. Both the egg quality and quantity from infected chickens is reduced due to infection. IBV infection with some strains of the virus prevents proper development of the reproductive tract of young chickens (4). Further, IBV infection can lead to immune suppression and an increase in secondary bacterial infections (5,6). Vaccines are available, however many limitations exist, such as adverse side effects in young chickens and rapid viral mutation leading to the emergence of new strains causing vaccines to become less effective (7).

Coronavirus genomes are approximately 30 kb in size, making them among the largest viral RNA genomes (8). To replicate their genomes, Coronaviruses encode 16 nonstructural proteins (nsps) in ORF1a and ORF1b in the first two-thirds of the genome (9). The latter third of the genome encodes structural proteins and virus-specific accessory proteins (9). Upon viral entry and genome release into the host cytoplasm, ORF1a/b are translated into polyproteins pp1a and pp1ab by host ribosomes (10). The polyproteins are cleaved by viral proteases to produce nsp subunits, which then remodel ER membranes to induce the formation of double membrane vesicles (DMVs), the sites of viral RNA replication (11-14). An important aspect of replication in this compartment is the production of a double-stranded RNA (dsRNA) intermediate, which if detected in the cytoplasm, triggers the host innate immune response (15). Host innate immune sensors that are triggered by dsRNA escape from the DMV during coronavirus infection include MDA5, PKR, and OAS/RNaseL, which induce robust production of type I and III IFN, PKR-mediated apoptosis, RNA degradation, and a reduced viral titer (16-20). Nonstructural protein 15 (nsp15), which is associated with the DMV, helps limit the detection of dsRNA by the host innate immune system (15). Nsp15 is an endoribonuclease which cleaves both ssRNA and dsRNA *in vitro* (9,21-23). In cells infected with viruses containing a catalytically knocked out nsp15, dsRNA accumulates and interferon responses increase compared to wild type (WT) virus infections (15,24-26). Nsp15 cleaves RNA 3’ of uridines, with a preference for a purine 3’ of the cleaved uridine and has been shown to explicitly target unpaired or flipped uridines (27-30). To carry out its enzymatic function, nsp15 performs a transesterification reaction using its active site catalytic triad containing two histidines and one lysine, akin to the RNaseA active site and indicative of a metal-independent cleavage mechanism (31-36). Briefly, nsp15 carries out a two-step acid-base reaction of transesterification and hydrolysis which results in a terminal 2’-3’ cyclic phosphate (18,37,38). For *in vitro* cleavage assays, nsp15 requires 5 mM manganese ions to carry out its enzymatic function, though this is believed to facilitate RNA binding not RNA cleavage directly (9,21-23). This concentration of manganese is not physiological and published cryo-electron microscopy (cryo-EM) and X-ray crystallography structures do not suggest the presence of metal ions in the active site (30,33,37,39-41). At least one recent study showed manganese is not required for nsp15 activity (37).

A nidovirus uridylate-specific endoribonuclease (NendoU/EndoU) homologue is encoded by members of multiple families within the *Nidovirales* order, including *Coronaviridae, Arteriviridae*, and *Tobaniviridae* (20,42-46). Conservation, based on amino acid sequence identity among coronavirus species across genera is 31-51%, with the active site catalytic triad having 100% sequence identity (**Supplementary Figure S1**) (32,47,48). Nsp15 has three recognized domains: the N-terminal domain (NTD), middle domain (MD), and C-terminal domain (CTD), residues 1-63, 64-171, and 196-338 in IBV nsp15, respectively, with a combined mass of 38 kDa (**Figure 1**). The NTD is responsible for oligomerization of the protein into a homohexameric assembly of 228 kDa which consists of a dimer of trimers (22,49). The MD is a variable domain and its function remains unknown (33). The endonuclease active site containing the catalytic triad is located within the nsp15 CTD (36).

**Figure 1.**
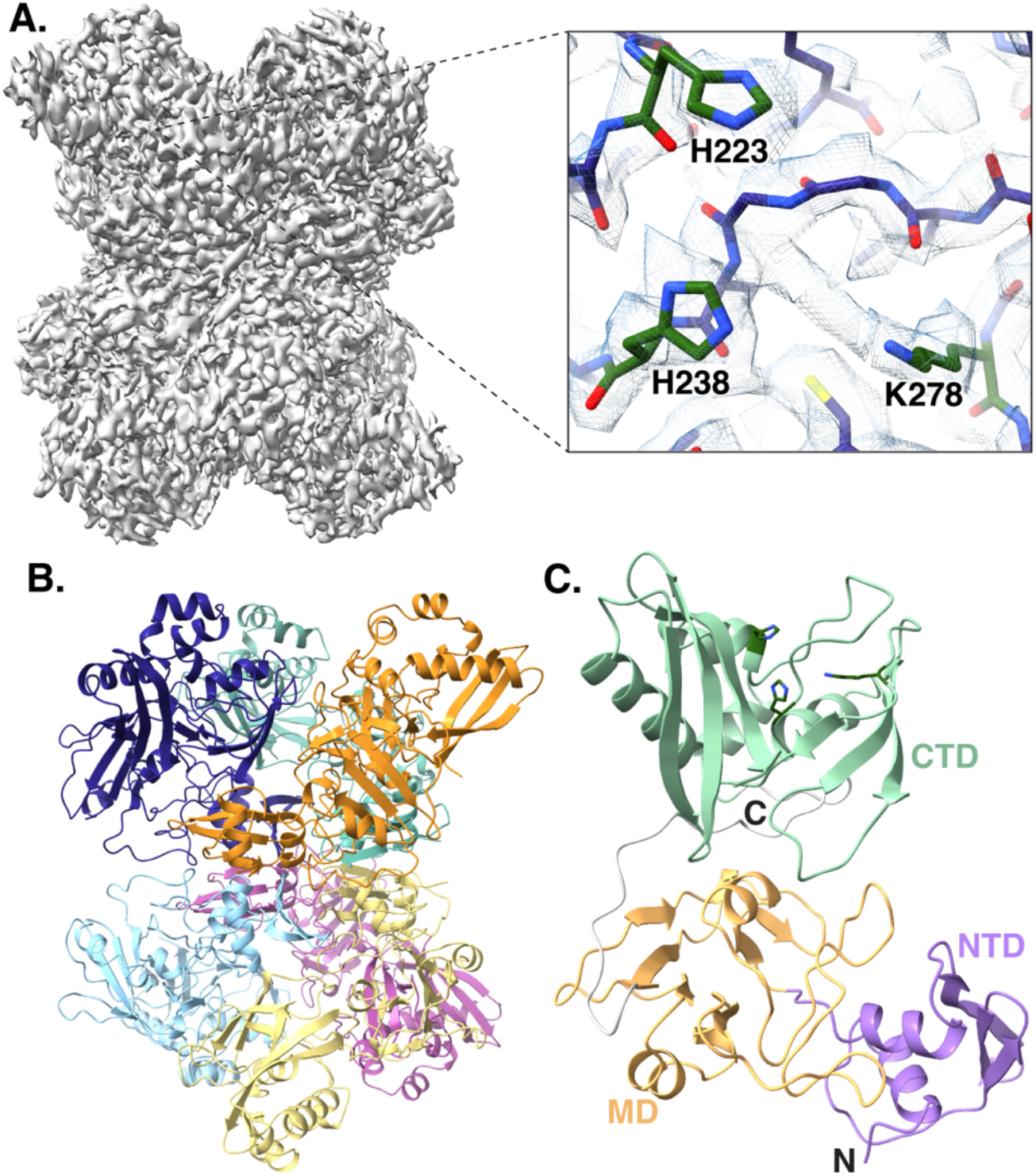
Cryo-EM structure and protein assembly of the apo-IBV nsp15. (**A**) 2.4 Å map of the apo IBV nsp15 and a zoomed in view of the catalytic triad with catalytic residues colored dark green. (**B**) Side view of the hexameric assembly of the apo-IBV nsp15, as a dimer of trimers. (**C**) The monomeric subunit domain organization: N-terminal domain (NTD, purple), middle domain (MD, orange), and the C-terminal endoribonuclease domain (CTD, green), with active site residues colored as in (A).

This study focuses on the *Gammacoronavirus* IBV nsp15. Conservation of nsp15 within the *Gammacoronavirus* genus, ranges from 41-95% sequence identify (**Supplementary Figure S1**). IBV’s genome is 27.4 kb and encodes 15 rather than 16 nsps as it lacks the nsp1 homologue found in other coronavirus genera (50). Despite extensive structural characterization of nsp15s from *Betacoronavirus* and some studies of nsp15 from *Alphacoronavirus*, nsp15 from *Gammacoronavirus* remains understudied (28,30,33,40,48). Here we present structures of the *Gammacoronavirus* IBV nsp15 apo-, single dsRNA-bound, and dual-dsRNA bound structures along with a biochemical characterization of nsp15 RNA cleavage activity. Our dsRNA-bound IBV nsp15 structure has many similarities to previously described *Betacoronavirus* dsRNA-bound SARS-CoV-2 nsp15 structures (30,51), however some of the residues peripheral to the active site differ between *Betacoronavirus* and *Gammacoronavirus*. Similar to SARS-CoV-2 nsp15 structures bound to dsRNA, we also notice that the RNA in our dsRNA-bound IBV nsp15 structure is not in perfect A-form, rather it is bent near the putative cleavage site. We hypothesize that IBV nsp15 accommodates a bent dsRNA to position a flipped uridine nucleotide into proximity of the catalytic triad for RNA cleavage and that variable active site peripheral residues which differ between SARS-CoV-2 and IBV nsp15 could alter RNA binding by IBV nsp15.

## MATERIALS AND METHODS

### Nsp15 construct design

The codon optimized IBV nsp15 sequence (Genscript, accession number: ACJ12832.1 5,972-6,309) was cloned into a pET46 bacterial expression vector containing an N-terminal 10X His-tag, a SUMO tag to enhance protein expression, a TEV protease cleavage site to remove tags and a GSGSAG linker to improve tag cleavage. Mutants were cloned using site-directed mutagenesis, with WT IBV nsp15-pET46 as the template DNA. The codon optimized SARS-CoV-2 nsp15 sequence (Genscript, accession number: QJA41639.1) was cloned into a pET46 bacterial expression vector containing an N-terminal 10X His-tag, a TEV protease cleavage site to remove tags and a GSGSAG linker to improve tag cleavage.

### Protein expression and purification

Constructs were transformed into C41 *E. coli* cells (Novagen), inoculated into 100 mL LB with 100 µg/mL ampicillin, and grown overnight at 37°C for approximately 16 hours. 10 mL from the 100 mL starter culture was used to inoculate 1 L of Terrific Broth with ampicillin and cells were grown at 37°C to an OD_600_ of 0.8-1.0. Before induction, cells were placed at 4°C for one hour and were occasionally swirled. Cooled cells were induced with 0.2 mM IPTG and placed at 16°C shaking overnight for approximately 20 hrs. Cells were pelleted at 3,500 x g for 15 minutes at 4°C and pellets were resuspended in 50 mL of lysis buffer 30 (50 mM Tris, 500 mM NaCl, 30 mM imidazole, 5% glycerol, and 2 mM DTT at pH 8) at room temperature. Cells were lysed in a LM20 microfluidizer (Microfluidics) at 20,000 psi and lysates were cleared by centrifugation at 25,000 x g for 30 minutes at 4°C. To remove nucleic acid, polyethyleneimine (PEI, Sigma P3143) was added to the supernatants at room temperature to a final concentration of 0.26% (w/v). PEI was added slowly over five equal additions every 5 minutes with the last incubation being 15 minutes. Precipitated nucleic acid was pelleted at 35,000 x g for 30 minutes at 20°C. Supernatant was retained and nsp15 protein was precipitated using 45 g of ammonium sulfate per 100 mL of supernatant. This solution was placed at 4°C overnight with slow stirring. Protein was pelleted at 30,000 x g for 30 minutes at 4°C. The supernatant was discarded and protein pellets were resuspended in 40 mL of lysis 30 buffer at room temperature. Resuspended pellets were spun at 30,000 x g for 30 minutes at 20°C to pellet aggregated protein. Supernatants were filtered (0.45 µm) and batch bound to 2 mL of Ni-NTA beads (Qiagen) for 30 minutes at room temperature. Beads were collected by centrifugation at 500 x g for 5 minutes at 20°C and loaded onto a column for washes and elution. Beads were washed with 20 mL of lysis 30 buffer (50 mM Tris, 500 mM NaCl, 30 mM imidazole, 5% glycerol, and 2 mM DTT at pH 8), followed by 20 mL of lysis 60 buffer (50 mM Tris, 500 mM NaCl, 60 mM imidazole, 5% glycerol, and 2 mM DTT at pH 8). Proteins were eluted with 10 mL of elution buffer (50 mM Tris, 500 mM NaCl, 500 mM imidazole, 5% glycerol, and 2 mM DTT at pH 8) and dialyzed overnight at room temperature in 1 L of dialysis buffer (50 mM Tris, 150 mM NaCl, 5% glycerol, 2 mM CaCl_2_, and 2 mM DTT at pH 8) with 2% (w/w) TEV protease in 3.5K MWCO SnakeSkin tubing (Pierce). Dialyzed and cleaved protein was added to 2 mL of Ni-NTA beads in a gravity column and incubated for 5 minutes. Unbound (cleaved) protein was collected and concentrated to 500 µL before spinning at 16,000 x g for 2 minutes and loading onto a Superdex200 Increase 10/30 GL size exclusion column. The size exclusion column was run with SEC buffer (50 mM Tris, 150 mM NaCl, 5% glycerol, 5 mM MnCl_2_ and 5 mM DTT at pH 7.5). The peak corresponding to hexameric protein was collected, concentrated in a 10K MWCO ultrafiltration concentrator (Amicon), flash frozen in liquid nitrogen, and placed at -80°C for storage.

### *In vitro* cleavage assay

IBV nsp15 WT and mutant protein activities were characterized using both single and double stranded RNA. 50 nM of purified protein was incubated with 250 nM of either a single-stranded 16 nucleotide (nt) RNA substrate with a single uridine cleavage site (6-FAM-GAAGCGAAACCCUAAG) or a double-stranded 35 nt RNA substrate with multiple uridine cleavage sites, modeled after the dsRNA substrate from Frazier et. al. 2022 (30) (6-FAM-UUUAGAUUUCAUCUAAACGAACAAACUAAAAUG UC with the complimentary strand GACAUUUUAGUUUGUUCGUUUAGAUGAAAUCUAAA). For experiments varying RNA and protein ratios, 250 nM RNA was kept constant and protein was used at 25, 50, 125, and 250 nM. The cleavage reactions were run in 20 mM Bis-tris, 150 mM sodium acetate at pH 6, 1 mM DTT, and 5 mM MnCl_2_ at 33°C. Samples were taken at 5, 10, 30, and 60 minutes and the reaction was stopped using a 2X RNA loading buffer (9.5 mL formamide, 20 mM EDTA, and 5 mg orange G). Samples were run on two-layer 8 M urea 10% and 20% PAGE for 70 minutes at 220 V. Gels were imaged with an Amersham Typhoon RGB and replicates were quantified in Fiji 2.16.0/1.54p (52).

### Cryo-EM sample preparation

For the dataset of nsp15 hexamer, purified IBV WT nsp15 at 1.5 mg/mL was spotted onto UltrAufoil R1.2/1.3 300 mesh grids (Quantifoil) and plunge frozen into liquid ethane. For the dsRNA-nsp15 complexed dataset, dsRNA was prepared at 50 µM in RNA binding buffer (10 mM KCl, 10 mM HEPES, 2 mM MgCl_2_ at pH 7.4) by heating at 75°C for 5 minutes and slow cooling for 90 minutes. The 35 nt dsRNA substrate used was the same as that used for dsRNA cleavage assays. Catalytically inactive IBV nsp15 H223A was complexed to dsRNA at a 1:1.2 molar ratio in 10 mM Bis-tris pH 6, 10 mM sodium acetate, 1 mM DTT, and 5 mM MnCl_2_ for 2 hours at 33°C. 3 µL of 1.5 mg/mL protein complexed to dsRNA was spotted onto UltrAufoil R1.2/1.3 300 mesh grids. Grids were blotted for four seconds in a Vitrobot Mark IV system (ThermoFisher) at 100% humidity, 15°C, and with a blot force of -8.

### Cryo-EM data collection and processing

For the apo nsp15 dataset, 4,323 movies were collected on a Talos Arctica (ThermoFisher) operating at 200 kV at 79,000x nominal magnification using a Gatan K3 direct electron detector. The average electron dose per movie was 60 e^-^/Å^2^ and the total exposure time per movie was 4.9 seconds. In CryoSPARC (53), movies were motion corrected using Patch Motion Correction and the defocus was estimated using Patch CTF Estimation. Using a minimum particle diameter of 100 Å and a maximum particle diameter of 130 Å, 4,029,726 particles were extracted for processing. Particles were sorted by 2D classification with 50 classes. A set of 3,434,267 particles was chosen for ab initio reconstruction, which was performed with two classes. One ab initio class containing 2,026,081 particles was selected for homogeneous refinement with D3 symmetry applied and had a final global resolution of 2.4 Å.

For the dsRNA-IBV nsp15 dataset, 10,990 movies were collected. In CryoSPARC, movies were motion corrected using Patch Motion Correction and the defocus was estimated using Patch CTF Estimation. Using a minimum particle diameter of 100 Å and a maximum particle diameter of 110 Å, the blob picker was used to identify 6,296,566 particles. Particles were classified three times using 2D classification with 50 classes. A set of 1,913,989 particles was chosen for ab initio reconstruction, which was performed once with three classes. A single class was used for reconstruction and homogeneous refinement with D3 symmetry applied. The refined particle orientations were symmetry expanded, to account for multiple dsRNAs bound to the same particle, and a focus mask was created which encompassed one protomer and the dsRNA density. The symmetry expanded particle stack was classified using a focus mask encompassing a single protomer and potentially bound RNA. From this classification, a single class was further 3D classified and a single class from the second classification was then locally refined to a final reconstruction with a resolution of 3.0 Å.

To isolate maps of IBV nsp15 with multiple dsRNAs bound, particles from three classes from the second round of focused classification above were merged and duplicate particles were removed. This particle stack of 375,281 projections was used for 3D classification with 10 classes. Of the resulting classes, two showed nsp15 bound to two dsRNAs on opposite trimers and two showed nsp15 bound to two dsRNAs on the same trimer. Equivalent poses were merged and locally refined to yield final maps containing 65,490 and 76,617 particles respectively, each at 3.3 Å resolution.

The IBV nsp15 hexameric protein coordinates were predicted using AlphaFold 3 (54) and fit into the apo map using ChimeraX-1.6 (55-57) and ISOLDE 1.12 (57). The coordinates were real-space refined and validated using Phenix-1.21.2-5419 (58) and minor structure adjustments were made in COOT 0.9.8.95 EL (ccp4) (59-61). The refined coordinates from the IBV nsp15 apo structure were used to fit into the singly dsRNA-bound map and dsRNA was modeled in COOT and refined using Phenix and ISOLDE. Despite reasonable global resolutions, the doubly dsRNA bound IBV nsp15 maps did not present high resolution density for the RNA for confident real-space refinement. The singly RNA-bound nsp15 protomer and nsp15 hexamer were used to rigid-body dock protomer coordinates into these maps using ChimeraX.

## RESULTS

### Cryo-EM structure of the apo-WT IBV nsp15

To compare nsp15 structure and activity across genera, we expressed and purified recombinant nsp15. By size exclusion chromatography, *Betacoronavirus* SARS-CoV-2 nsp15 elutes as hexamer and monomer peaks (**Supplementary Figure S2**) (30). However, *Gammacoronavirus* IBV nsp15 elutes as primarily a hexamer, with a minimal monomer peak (**Supplementary Figure S2**). We confirmed the activity of IBV nsp15 WT and nsp15 H223A catalytic knockout proteins using a gel-based in vitro RNA cleavage assay (**Supplementary Figure S2**).

We used single particle cryo-electron microscopy (cryo-EM) to characterize the IBV nsp15 structure. The first cryo-EM dataset used the IBV nsp15 hexameric protein alone. The 2D class averages show many representations of the IBV nsp15, such as top and side views (**Supplementary Figure S3, S4**). A 3D map was refined using D3 symmetry to produce a 2.4 Å structure of the apo-*Gammacoronavirus* IBV nsp15 enabling coordinate modeling (**Figure 1 and Table 1**). The nsp15 homohexamer reveals structural conservation across the coronavirus genera with NTD, MD and CTD domains for each nsp15 protomer and conserved oligomerization protomer interfaces to form a dimer of trimers (**Supplementary Figure S1**). The active site catalytic triad architecture is maintained with the same positioning as those of the *Betacoronavirus* MHV, MERS-CoV, SARS-CoV, and SARS-CoV-2 nsp15s, as well as the *Alphacoronavirus* 229E nsp15.

### Cryo-EM structure of the dsRNA-bound IBV nsp15

To examine IBV nsp15’s RNA binding mechanism, we prepared a cryo-EM sample using catalytically inactive nsp15 H223A bound to a 35 nt dsRNA with multiple potential uridine substate cleavage sites. The substrate nucleotide sequence was based on a previously described substrate used for dsRNA-bound SARS-CoV-2 nsp15 (30). 10,990 movies were collected from two combined datasets. 2D classes revealed the presence of bound dsRNA, represented by lighter grey whisps extending from the hexameric nsp15 (53). After ab initio reconstruction, volumes were refined using homogenous refinement with D3 symmetry. Particle sets from homogenously refined volumes were symmetry expanded and a mask encompassing a single nsp15 protomer and putative RNA was applied during 3D classification, identifying all nsp15 protomers in the dataset bound to RNA. The final dataset contained 180,637 protomer projections and was locally refined, resulting in a final map of 3.1 Å (**Figure 2, Supplementary figure S5 and S6, and Table 1**). This map allowed coordinate modeling of the hexameric protein as well as 22 nt of RNA in one strand and 20 nt of RNA in the complimentary strand. Given the possibility of multiple RNA binding poses to place a uridine in the nsp15 active site, the density for the RNA does not distinguish nucleotide bases despite resolved base and backbone map density. An RNA sequence is modeled representing one possible pose.

**Figure 2.**
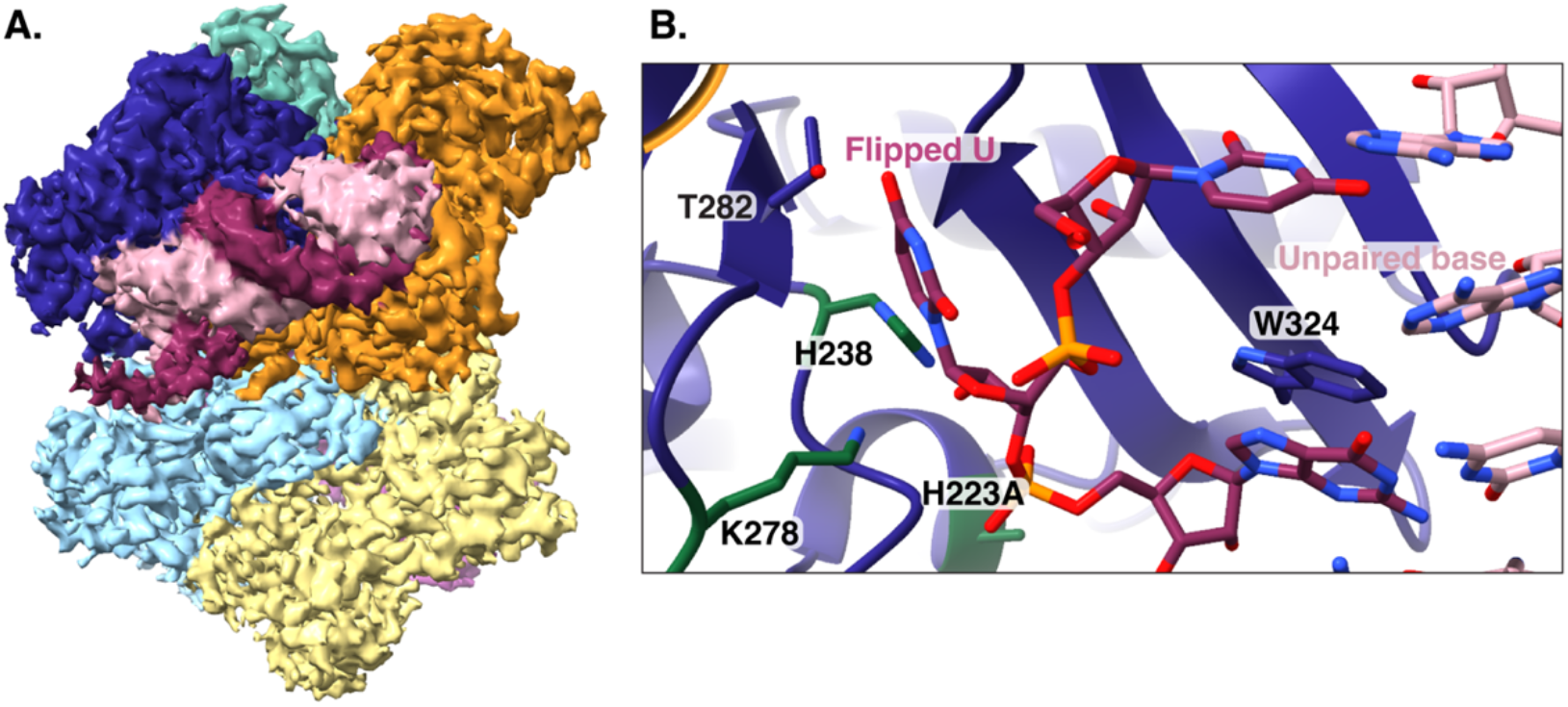
Cryo-EM structure of the dsRNA-bound IBV nsp15. (**A**) 3.0 Å isosurface map of dsRNA bound to IBV nsp15. (**B**) Zoomed in view of the flipped uridine in proximity to the active site.

Similar to the structure of SARS-CoV-2 nsp15 bound to dsRNA, the IBV nsp15 engages a uridine nucleotide that is flipped out of the dsRNA helix (**Figure 2**). Recognition of the uridine base occurs by hydrogen bonding to IBV nsp15 peripheral residue T282 main and side chains while catalytic residues H238 and K278 hydrogen-bond to the uridine ribose 2’ hydroxyl. Protein interactions at the uridine 2’ hydroxyl may be important for ribose deprotonation and formation of a 2’-3’ cyclic phosphate product. The nucleotide following the flipped-out uridine has been modeled as a guanosine and stacks on W324 potentially stabilizing the non-A-form structure. It has previously been hypothesized that SARS-CoV-2 nsp15 uses a base-flipping mechanism to engage the uridine to be cleaved (30). However, a later paper, suggested that a uridine can flip out from the double-stranded helix spontaneously and nsp15 takes advantage of this process for substrate recognition (29).

### IBV nsp15 utilizes peripheral residues to engage dsRNA

Our structure with dsRNA density reveals the dsRNA is bent to interact with nsp15 and is not in a perfect A-form confirmation. While the residue following the recognized uridine forms base-pairing interactions, the uridine itself does not, leaving an unpaired adenosine in the uncleaved strand (**Figure 2**). The flipped-out uridine and unpaired base create a significant distortion in the RNA helix causing a bend. To accommodate the RNA bend, IBV nsp15 uses residues peripheral to the active site. Despite the observation of bent RNA with both SARS-CoV-2 and IBV nsp15 structures, peripheral RNA interacting residues differ between *Betacoronavirus* and *Gammacoronavirus* nsp15s (30).

We selected several IBV nsp15 residues that we observed to interact with the dsRNA for further study by mutagenesis and biochemical assays (**Figure 3**). Nearest the active site and the flipped-out uridine are K306 and S307, which bind the RNA in the RNA minor groove. K306 is similarly a lysine in porcine deltacoronavirus (PDCV) nsp15 but a valine in 229E and SARS-CoV-2 nsp15. The positive charge of K306 could have significant interactions with the negatively charged dsRNA backbone. S307 is conserved across coronavirus genera and spatially resides beneath W324, which supports the purine 3’ of the flipped-out uridine nucleotide. S307 may also hydrogen bond with the non-cleaved strand. 3’ of the putative cleavage site in the RNA major groove, a short IBV nsp15 loop (amino acids 227-236) contacts the RNA backbone. At the apex of this loop near the RNA is IBV nsp15 D230, K231, and P232. Nsp15 D230 is a modestly conserved serine across coronavirus genera with the exception of *Alphacoronavirus* members where this position is a conserved serine. Going from a hydrophilic hydroxyl to a negatively charged carboxyl in IBV may cause repulsion with the nearby RNA backbone in IBV nsp15, weakening RNA binding compared to other coronavirus members. *Alphacoronavirus* 229E nsp15 also has a lysine at the IBV K231 equivalent position but this residue is a histidine in *Betacoronavirus* SARS-CoV-2 nsp15 and a serine in *Deltacoronavirus* PDCV nsp15. IBV nsp15 P232 is also a proline in PDCV nsp15 but is a serine in SARS-CoV-2 and a threonine in 229E nsp15. Nearly a full RNA helical turn away from the flipped-out uridine, Q138 of an adjacent nsp15 protomer contacts the RNA minor groove, primarily interacting with RNA bases. Q138 is not conserved across the coronavirus genera (glycine in SARS-CoV-2 and threonine in 229E and PCDV nsp15).

**Figure 3.**
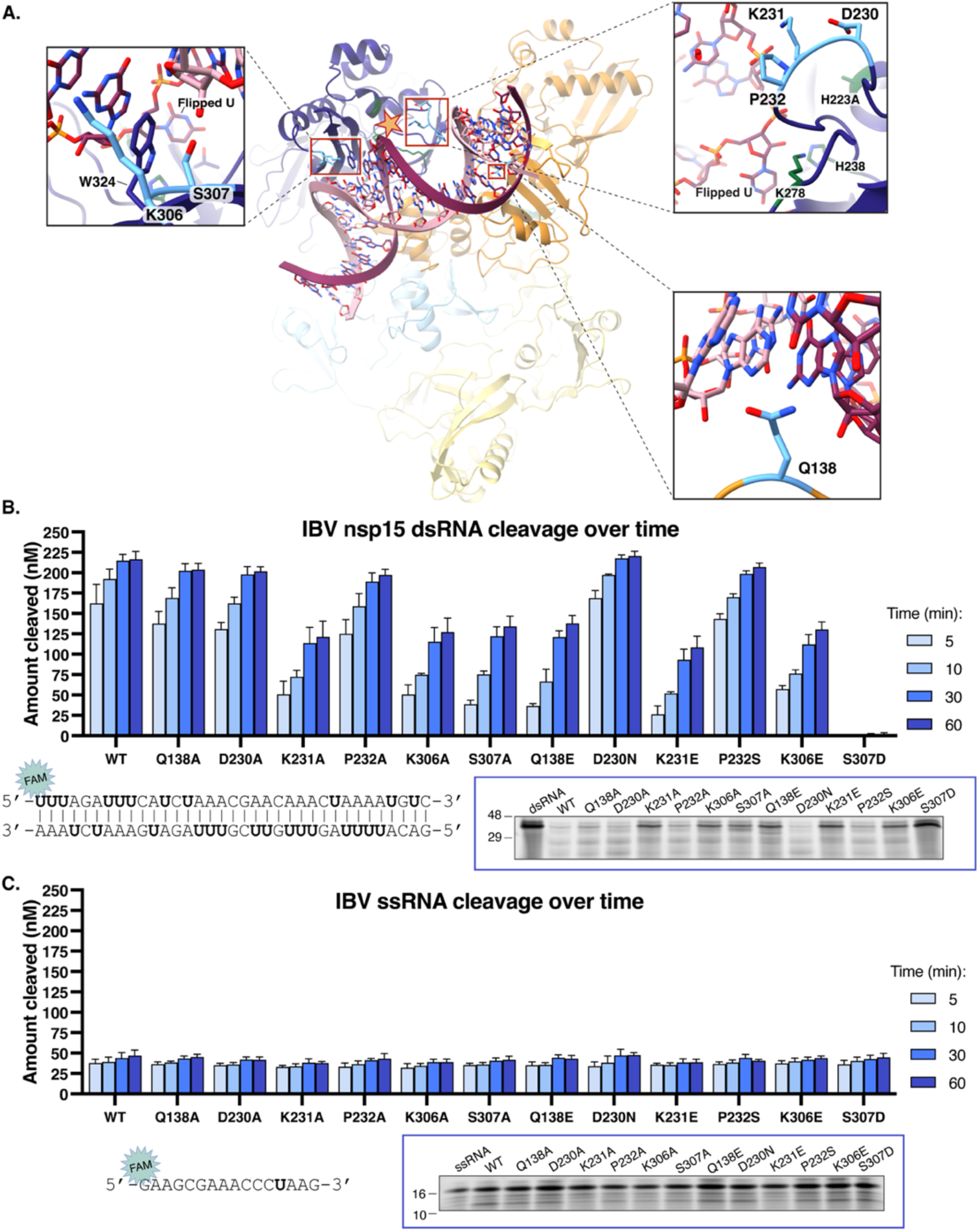
Peripheral active site residues confer IBV nsp15 substrate specificity. (**A**) The coordinate model of IBV nsp15-dsRNA shows several protein positions peripheral to the active site that contact the dsRNA. Residues chosen for mutation are colored in blue and active site residues in dark green are also indicated by a star. Nsp15 wild type and mutant proteins cleave dsRNA (**B**) and ssRNA (**C**). The nucleic acid substrates each contained a single 5’ 6-FAM fluorescent tag and sequences are shown below the graphs, along with a representative gel of a 60-minute RNA cleavage reaction. Graphs in (B) and (C) represent three independent replicates. See also Fig. S8 and S9.

Active site peripheral residues in IBV nsp15 that we hypothesized as important for dsRNA binding were singly mutated and then recombinantly expressed. Size-exclusion chromatography suggested all mutant proteins form hexameric oligomers (**Supplementary figure S7**). The wild type and mutant proteins (50 nM) were used in gel-based *in vitro* cleavage assays with 250nM of dsRNA or ssRNA. IBV nsp15 mutants Q138A, D230A, and P232A have dsRNA cleavage activity similar to WT. IBV nsp15 mutants D230N and P232S have modestly higher dsRNA cleavage activity compared to WT. K231A, K306A, S307A, Q138E, K231E, and K306E mutants have less dsRNA cleavage activity compared to WT and nsp15 S307D has no dsRNA cleavage activity observed (**Figure 3, Supplementary Figure S8, S9**).

W324 helps stabilize the nucleotide base 3’ of the flipped uridine to support the scissile phosphate in the active site (30). S307 sits just beneath W324 and may both provide van der Waals interactions to position the W324 side chain and hydrogen-bonding to the uncleaved RNA strand, suggested by a significant loss of activity for S307A (**Figure 3**). Mutation to S307D introduces a negative charge that may repel the RNA backbone of the uncleaved strand while also potentially disrupting the conformation of W324, leading to a loss of the pi-stacking interaction with nucleotide bases. K306 may help to position the ribose sugar of the nucleotide base-pairing to the W324-stacked purine, stabilizing the flipped-out uridine conformation (30). This interaction would be lost in nsp15 K306A, and nsp15 K306E may create unfavorable electrostatics with the nearby phosphate backbone. K231 contacts the negatively charged RNA backbone of the cleaved strand. K231A and K231E would lose this interaction possibly leading to their reduced activity. D230 may partially neutralize the K231 positive charge. Mutation to D230N may prevent charge neutralization without otherwise perturbing the loop leading to better substrate binding and the observed modest increase in dsRNA cleavage activity. In the same loop, P232S may provide a more hydrophilic surface for binding the uncleaved RNA strand. Nsp15 dsRNA cleavage reduction for Q138E indicates that at least two distinct protomers within the hexamer can impact the observed cleavage activity. Because Q138A has activity equivalent to wild type nsp15, the second protomer likely accommodates the dsRNA rather than being required for the observed cleavage.

### IBV nsp15 preferentially cleaves dsRNA over ssRNA

Despite prior reports of ssRNA cleavage by coronavirus nsp15, we observe minimal, though detectable, ssRNA cleavage by wild type IBV nsp15 (**Figure 3**). In contrast, dsRNA substrates were cleaved much more efficiently. To evaluate IBV nsp15’s cleavage ability of dsRNA versus ssRNA, increasing amounts of WT IBV nsp15 (25, 50, 125, and 250 nM) were incubated with 250 nM of RNA (**Figure 4**). Our data shows that at the same protein concentrations, more dsRNA is cleaved compared to ssRNA. This aligns with SARS-CoV and 229E nsp15 findings where dsRNA is more efficiently cleaved than ssRNA (9) and contrasts findings with SARS-CoV-2 nsp15 where ssRNA is cleaved more efficiently than dsRNA (30). A caveat to this observation is that the dsRNA substrate contains eleven uridine cleavage sites in the labeled strand compared to the single uridine in the ssRNA substrate. However, 50 nM WT IBV nsp15 cleaves ∼70% of the 250 nM dsRNA substrate by five minutes, yet only cleaves ∼30% of the ssRNA substrate in 60 minutes (**Figure 4**).

**Figure 4.**
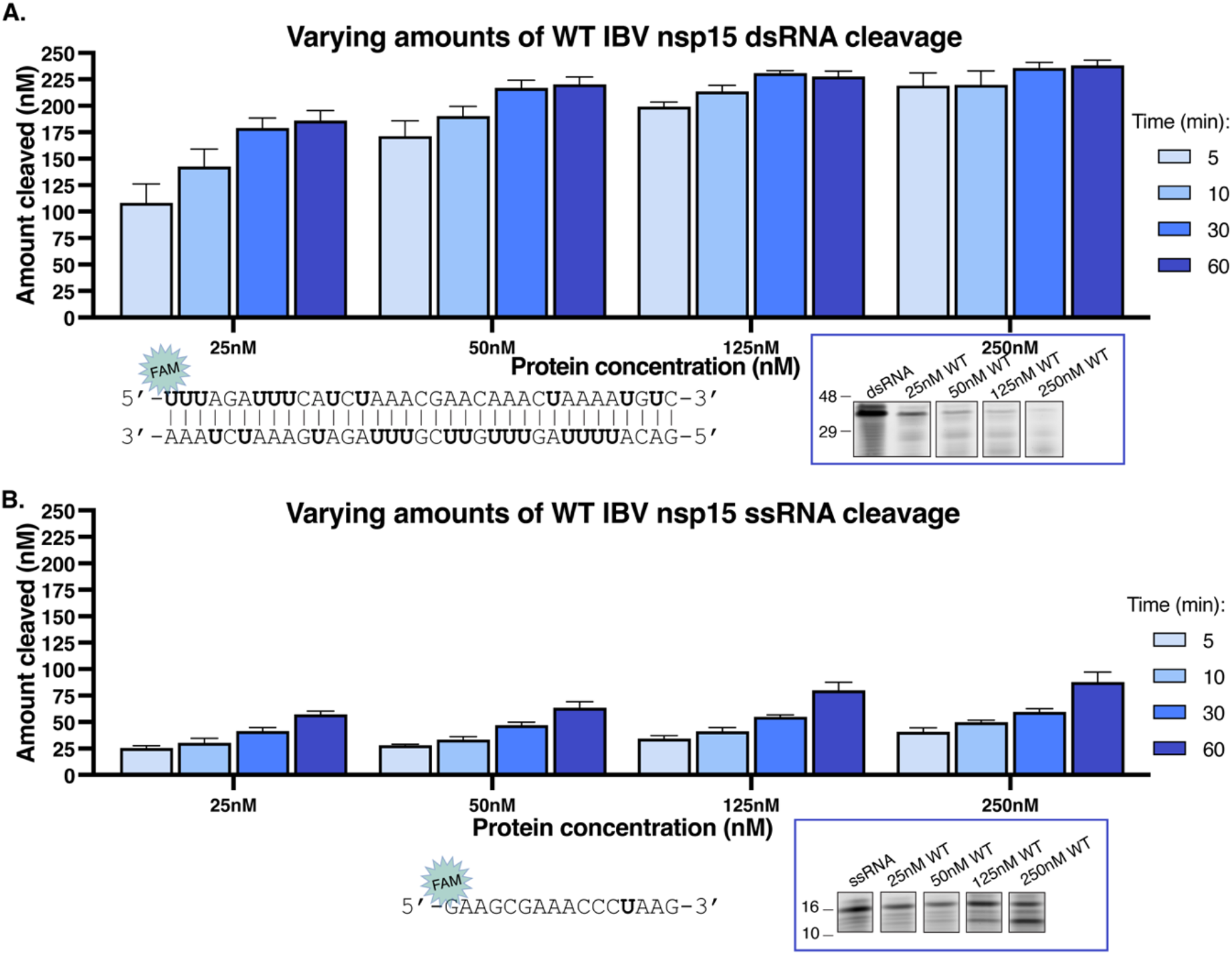
IBV nsp15 cleaves dsRNA preferentially over ssRNA. (**A**) Increasing concentrations of IBV nsp15 demonstrate efficient cleavage of a dsRNA substrate. (**B**) ssRNA is cleaved slowly by IBV nsp15. Graphs in (A) and (B) represent three independent replicates. See also Fig. S10 and S11.

### IBV nsp15 binds more than one dsRNA simultaneously

Further analysis of the dsRNA-IBV nsp15 cryo-EM dataset identified particle subsets containing two dsRNAs bound to IBV nsp15 resulting in two additional maps each with a global resolution of 3.3 Å (**Figure 5, Supplementary figure S5 and S12, and Table 1**). The local resolution of the dsRNA was lower, ∼5-7 Å. As these maps were of lower resolution, dsRNA-bound nsp15 coordinate models were rigid body fit into these maps without further refinement. Previously, researchers have speculated nsp15 may bind multiple dsRNAs simultaneously (30,33) and one group has determined a structure of a hexameric SARS-CoV-2 nsp15 bound to two dsRNAs where the dsRNAs engage two nsp15 protomers within a single trimer (62). One of our reconstructed maps of IBV nsp15 bound to two dsRNAs recapitulates the previously determined SARS-CoV-2 nsp15 structure with two dsRNAs bound at two protomers of the same trimer (**Figure 5A and 5B**). The second reconstructed map shows the two dsRNAs binding to nsp15 protomers on opposite trimers though the two engaged nsp15 active sites are not closely neighboring (**Figure 5C and 5D**). In the hexameric nsp15, the active site is surface-exposed presenting six possible active sites available for engaging multiple dsRNAs simultaneously. However, two dsRNAs binding at neighboring nsp15 active sites between trimers is anticipated to create clashes. Such clashes may be avoided by the inherent flexibility of RNA though such a binding pose has not yet been experimentally reported.

**Figure 5.**
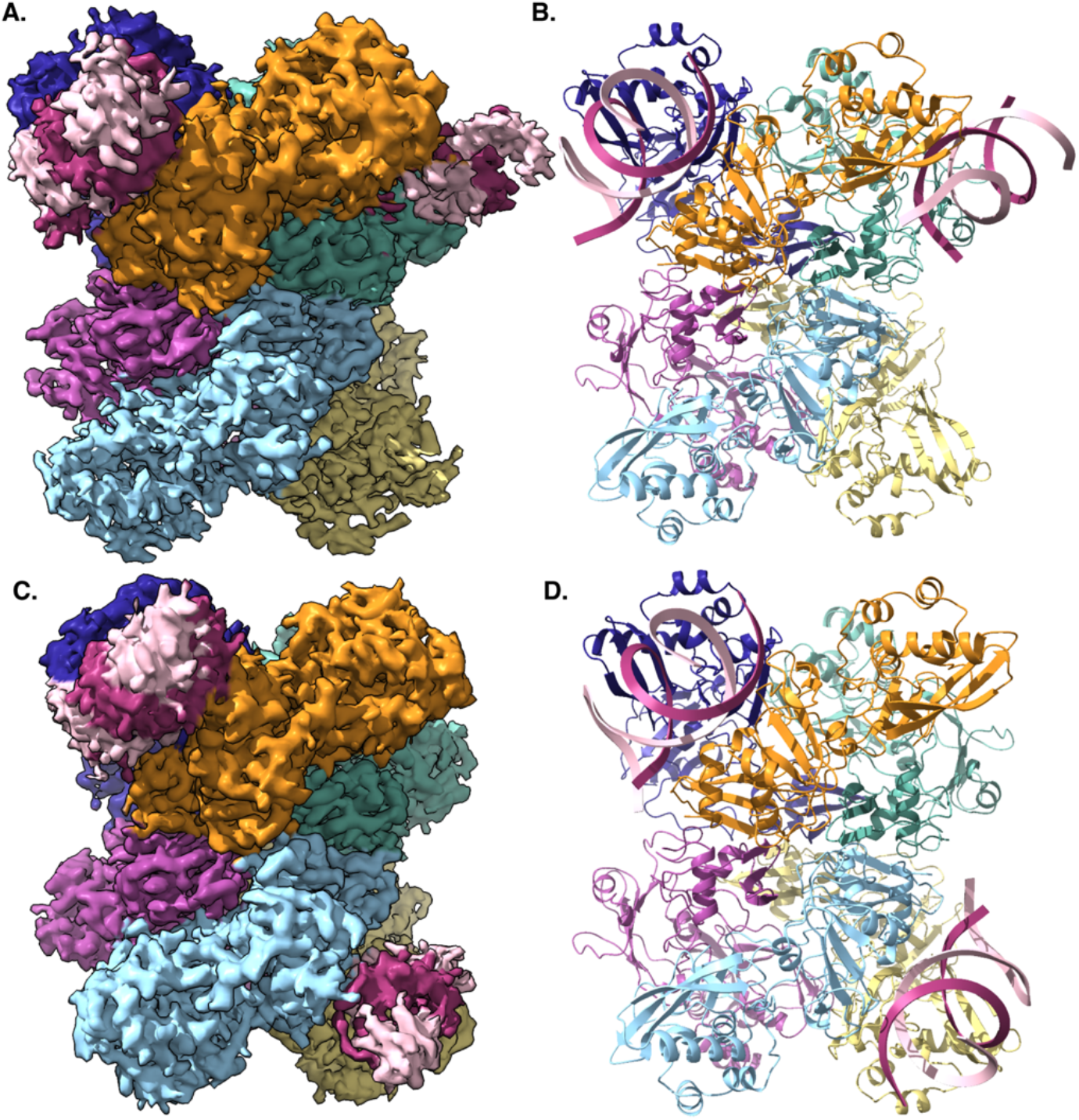
IBV nsp15 binds two dsRNAs simultaneously. Map (**A**) and model (**B**) of dual dsRNA-bound IBV nsp15 within the same trimer. Map (**C**) and model (**D**) of dual dsRNA-bound IBV nsp15 across two trimers.

## Discussion

The experiments presented here identify active site peripheral residues as being important for coronavirus nsp15 endoribonuclease activity. Our data clearly show a preference for IBV nsp15 to cleave dsRNA over ssRNA substrates. We also observe IBV nsp15 hexamers with two dsRNA substrates bound simultaneously.

Prior work has emphasized the importance of the active-site adjacent nsp15 W324 in pi-stacking with the nucleotide base 3’ of the uridine in the cleavage site. Here, we add, K306 and S307, creating a cluster of IBV nsp15 residues important for binding and positioning the scissile phosphate for cleavage. While species of the *Igacovirus* subgenus (*Gammacoronavirus Igacovirus*) like IBV and *Deltacoronavirus* PDCV have nsp15 K306, *Betacoronavirus* and *Alphacoronavirus* nsp15 have a mostly conserved valine at this position suggesting that the importance of this cluster may be specific to particular viral species. As noted in the results, the apex of the IBV nsp15 227-236 loop contacts dsRNA in the major groove, 3’ of the uridine RNA cleavage site. The positively charged K231 contacts the phosphate backbone, securing the dsRNA in place for cleavage of the flipped uridine. We saw less activity for both K231A and K231E nsp15 mutants with dsRNA compared to WT, whereas the analogous point mutation in SARS-CoV-2 nsp15, H243A, resulted in minimal defects (30). As we see large activity defects with the single point mutations in IBV nsp15, K231 perhaps has a more significant role in IBV ns15’s interaction with dsRNA than the analogous residue in SARS-CoV-2 nsp15.

IBV nsp15 has a strong propensity for cleavage of dsRNA over ssRNA *in vitro*. Only at high concentrations of WT IBV nsp15 and after a significant amount of time, do we observe IBV nsp15 cleavage of ssRNA. This result is in agreement SARS-CoV and 229E nsp15 cleavage preferences yet contrasts with SARS-CoV-2 nsp15 (9,21,30). dsRNA is hypothesized to be the physiological substrate of nsp15 during viral infection as the virus works to replicate undetected from the immune system.

The cryoEM maps revealing nsp15 with two bound dsRNAs support a previously described positive cooperative binding mechanism and allosteric sigmoidal RNA cleavage kinetics though we note no large differences between RNA-bound and unbound nsp15 RNA-binding sites (41). Binding more than one dsRNA at the same time could be how nsp15 is able to carry out allosteric sigmoidal kinetics. This also reintroduces the questions of what advantage there is for the virus to present six functional active sites, whether these sites are used simultaneously during infection, and how nsp15 oligomers might be incorporated into a holo-viral replication complex.

The conservation of the NendoU endonuclease across many species of *Nidovirales* indicates the importance of its role in subverting immune responses that are triggered by an accumulation of dsRNA. Combining structural observations with mutagenesis and *in vitro* biochemistry of IBV nsp15, we have demonstrated the impact that active site peripheral residues can have on IBV nsp15 dsRNA binding and cleavage. The results clearly show diminished or altered activity when these sites are mutated. Despite the impact that these residues have on enzyme activity, many of these residues show poor conservation across or even within coronavirus genera. The heterogeneity in active site peripheral residues and the impact these amino acids can have on nsp15 dsRNA cleavage indicates the existence of multiple solutions for nsp15 to engage RNA. This presents the possibility of evolutionary shaping of nsp15 activity using residue positions beyond the active site to tune endoribonuclease activity. Future work should focus on delineating the impacts of active site peripheral residues on RNA binding and catalysis with complementary viral reverse genetics experiments.

## SUPPLEMENTARY DATA

Supplementary Data are available at NAR online.

## ACKNOWLEDGEMENTS

A special thanks to Dr. Kelly Watters for her advice and experimental assistance and Dr. Susan C. Baker of Loyola University, Stritch School of Medicine for advice and support. Some of this work was performed in the Cryo-EM Research Center (CEMRC) in the Department of Biochemistry at the University of Wisconsin-Madison.

## AUTHOR CONTRIBUTIONS

Ena S. Tully: Conceptualization, Data curation, Formal analysis, Investigation, Methodology, Project administration, Validation, Visualization, Writing— original draft Robert N. Kirchdoerfer: Conceptualization, Formal analysis, Funding acquisition, Methodology, Project administration, Resources, Supervision, Validation, Visualization, Writing—review & editing

## FUNDING

This work was supported by fellowship from the University of Wisconsin-Madison Department of Biochemistry to E.S.T.; and the National Institute of Allergy and Infectious Diseases (NIAID) AI159945 to R.N.K. Funding for open access charge: UW-Madison, Department of Biochemistry.

## DATA AVAILABILITY

Cryo-EM structures can be accessed in the PDB and EMDB with the following accession codes: PDB: 37IX, EMDB-78219, PDB: 37KX, EMDB-78244, PDB: 37UN, EMDB-78493, PDB:37UO, EMDB-78494. All additional data displayed in this manuscript are available in the main text and the online supplementary material.

## Notes

https://www.rcsb.org/structure/unreleased/37IX

https://www.rcsb.org/structure/unreleased/37KX

https://www.rcsb.org/structure/unreleased/37UN

https://www.rcsb.org/structure/unreleased/37UO

### Competing Interest Statement

The authors have declared no competing interest.

