## Supplementary data for "Structural and biochemical analysis of the IBV nsp15 endoribonuclease reveals the necessity of peripheral site residues for activity"

| | Apo IBV nsp15 (EMDB-78219) (PDB 37IX) | dsRNA-bound IBV nsp15 (EMDB-78244) (PDB 37KX) | Dual dsRNA-bound IBV nsp15 (opposite trimers) (EMDB-78493) (PDB 37UN) | Dual dsRNA-bound IBV nsp15 (same trimer) (EMDB-78494) (PDB 37UO) |
|---|---|---|---|---|
| **Data collection and processing** | | | | |
| Microscope | Talos Arctica | Talos Arctica | | |
| Voltage (kV) | 200 | 200 | | |
| Detector | Gatan K3 | Gatan K3 | | |
| Dose Rate ($e^{-}$/pixel/sec) | 13.9 | 13.9 | | |
| Exposure Time (sec) | 4.9 | 4.88 | | |
| Electron exposure (e–/Å$^{2}$) | 60 | 60 | | |
| Frames (no.) | 60 | 60 | | |
| Defocus range (µm) | -0.7 to -1.7 | -0.7 to -1.7 | | |
| Data Collection Mode | EFTEM, Counting, CDS | EFTEM, Counting, CDS | | |
| Magnification | 79,000x | 79,000x | | |
| Pixel size (Å) | 1.063 | 1.064 | | |
| Movies Collected (no.) | 4,323 | 10,990 | | |
| Initial Particle Images (no.) | 4,029,726 | 6,296,566 | | |
| Final Particle Images (no.) | 2,026,081 | 180,637 | 65,490 | 76,617 |
| Symmetry imposed | D3 | C1 | C1 | C1 |
| Sphericity | 0.976 | 0.975 | 0.890 | 0.927 |
| Map resolution (Å) - GSFSC | 2.4 | 3.0 | 3.3 | 3.3 |
| **Model Refinement** | | | | |
| Initial model used (PDB code) | AlphaFold3 model | 37XI | 37KX | 37KX |
| Model composition | | | | |
| Non-Hydrogen Atoms | 16,254 | 17,502 | 17,694 | 17,639 |
| Protein Residues | 2,040 | 2,034 | 2,035 | 2,033 |
| Nucleic Acid Residues | 0 | 42 | 72 | 70 |
| R.M.S. deviations | | | | |
| Bond lengths (Å) | 0.014 | 0.003 | 0.004 | 0.004 |
| Bond angles (°) | 1.875 | 0.571 | 0.653 | 0.647 |
| Validation | | | | |
| MolProbity score | 0.50 | 0.87 | 0.94 | 0.93 |
| Clashscore | 0.00 | 1.37 | 1.80 | 1.72 |
| Rotamer outliers (%) | 0.00 | 0.00 | 0.00 | 0.00 |
| Ramachandran plot | | | | |
| Favored (%) | 98.5 | 98.5 | 98.5 | 98.5 |
| Allowed (%) | 1.5 | 1.5 | 1.5 | 1.5 |
| Outliers (%) | 0.0 | 0.00 | 0.00 | 0.00 |
| EMRinger Score | 5.0 | 4.2 | 2.9 | 2.7 |
| Q Score | 0.65 | 0.55 | 0.46 | 0.44 |

**Table S1.** Cryo-EM data collection, refinement and validation statistics (1-5).

**A.** Nsp15 Percent Identity Matrix

| | HKU11 | PDCV | FCoV | 229E | PEDV | HKU8 | HKU9 | MHV | HKU1 | SARS | SARS2 | HKU4 | MERS | HKU5 | SW1 | IBV | TCoV |
|---|---|---|---|---|---|---|---|---|---|---|---|---|---|---|---|---|---|
| HKU11 | 100 | 80 | 33 | 34 | 35 | 36 | 37 | 30 | 30 | 38 | 37 | 36 | 33 | 37 | 31 | 36 | 36 |
| PDCV | 80 | 100 | 35 | 36 | 35 | 36 | 37 | 32 | 30 | 37 | 38 | 37 | 35 | 39 | 32 | 39 | 38 |
| FCoV | 33 | 34 | 100 | 64 | 68 | 65 | 40 | 43 | 45 | 40 | 41 | 45 | 45 | 45 | 40 | 38 | 38 |
| 229E | 34 | 36 | 64 | 100 | 72 | 69 | 41 | 43 | 42 | 44 | 44 | 46 | 47 | 46 | 41 | 40 | 40 |
| PEDV | 35 | 35 | 68 | 72 | 100 | 72 | 45 | 45 | 46 | 45 | 45 | 49 | 51 | 50 | 40 | 42 | 42 |
| HKU8 | 36 | 36 | 65 | 69 | 72 | 100 | 45 | 45 | 45 | 45 | 46 | 48 | 49 | 48 | 37 | 39 | 41 |
| HKU9 | 37 | 37 | 40 | 41 | 45 | 45 | 100 | 44 | 44 | 48 | 47 | 46 | 43 | 46 | 36 | 41 | 40 |
| MHV | 30 | 32 | 43 | 43 | 45 | 45 | 44 | 100 | 76 | 49 | 48 | 48 | 48 | 47 | 35 | 36 | 36 |
| HKU1 | 30 | 30 | 45 | 42 | 46 | 45 | 44 | 76 | 100 | 49 | 50 | 46 | 47 | 45 | 35 | 38 | 37 |
| SARS | 38 | 37 | 40 | 44 | 45 | 45 | 48 | 49 | 49 | 100 | 89 | 50 | 50 | 51 | 36 | 40 | 40 |
| SARS2 | 37 | 38 | 41 | 44 | 45 | 46 | 47 | 48 | 50 | 89 | 100 | 51 | 51 | 52 | 36 | 39 | 39 |
| HKU4 | 36 | 37 | 45 | 46 | 49 | 48 | 46 | 48 | 46 | 50 | 51 | 100 | 77 | 81 | 33 | 39 | 39 |
| MERS | 33 | 35 | 45 | 47 | 51 | 49 | 43 | 48 | 47 | 50 | 51 | 77 | 100 | 82 | 36 | 37 | 38 |
| HKU5 | 37 | 39 | 45 | 46 | 50 | 48 | 46 | 47 | 45 | 51 | 52 | 81 | 82 | 100 | 35 | 33 | 38 |
| SW1 | 31 | 32 | 40 | 41 | 40 | 37 | 36 | 35 | 35 | 36 | 36 | 34 | 37 | 35 | 100 | 42 | 41 |
| IBV | 36 | 39 | 38 | 40 | 42 | 39 | 41 | 36 | 38 | 40 | 39 | 38 | 37 | 37 | 42 | 100 | 95 |
| TCoV | 36 | 38 | 38 | 40 | 42 | 41 | 40 | 36 | 37 | 40 | 39 | 39 | 38 | 38 | 41 | 95 | 100 |

Least similar — Most similar

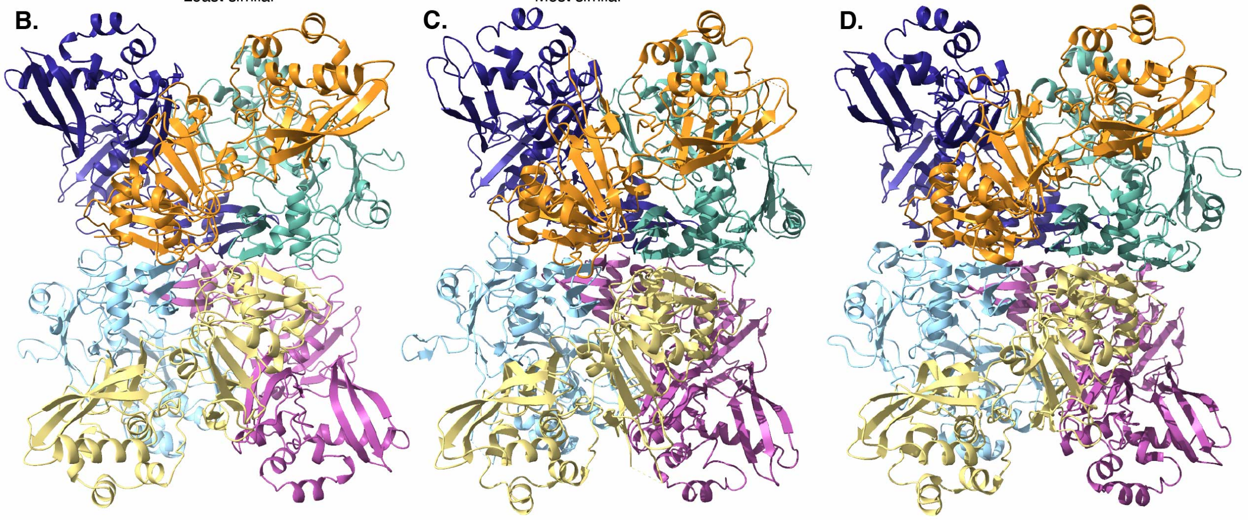


IBV nsp15: 37IX

229E nsp15: 4RS4

SARS-CoV-2 nsp15: 7N33

**Supplementary Figure 1**. Sequence and structural similarity across coronavirus nsp15 genera. (**A**) Percent identity matrix of nsp15 from various genera. HKU11 and PDCV are from the *Deltacoronavirus* genus. FCoV, 229E, PEDV, and HKU8 are from the *Alphacoronavirus* genus. HKU9, MHV, HKU1, SARS-CoV, SARS-CoV-2, HKU4, MERS-CoV, and HKU5 are from the *Betacoronavirus* genus. SW1, IBV, and TCoV are from the *Gammacoronavirus* genus. Comparison of (**B**) the *Gammacoronavirus* IBV nsp15 structure from this study with representative nsp15s from the (**C**) *Alphacoronavirus* genus and (**D**) *Betacoronavirus* genus display structural similarity across the coronavirus genera.

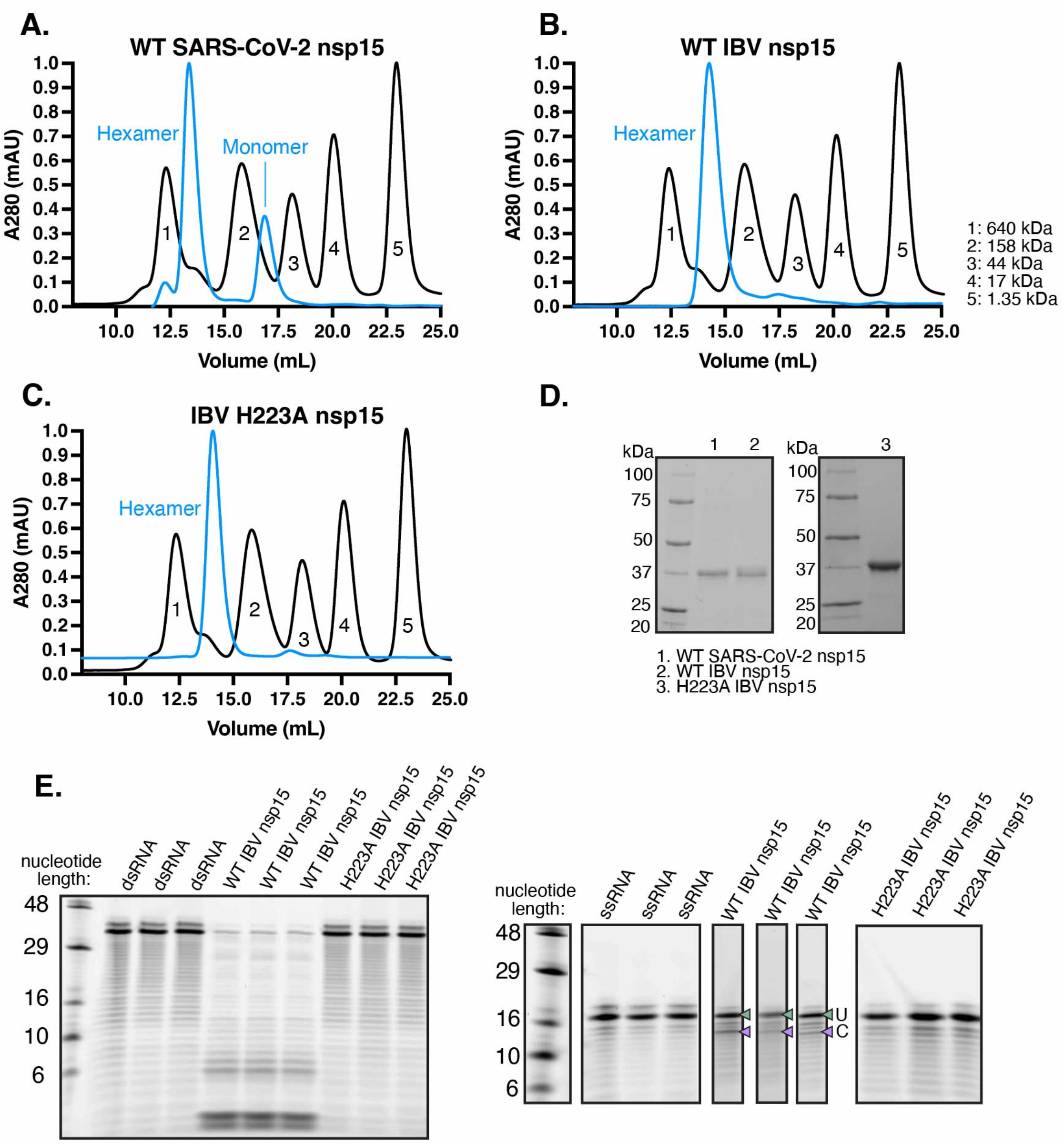


**Supplementary figure 2.** Purification of control nsp15s. (**A**) The SEC profile of WT SARS-CoV-2 nsp15 shows a hexamer and monomer peak, whereas SEC profiles of WT IBV nsp15 (**B**) and H223A IBV nsp15 (**C**) show just one main peak for hexamer protein. (**D**) SDS-PAGE of purified WT SARS-CoV-2, WT IBV nsp15, and H223A IBV nsp15 post-SEC. (**E**) dsRNA (left) and ssRNA (right) cleavage assay with 50 nm H223A IBV nsp15 after 60 min show H223A has no activity (U: uncleaved band and C: cleaved band). Cleavage reactions are presented as three technical replicates for each condition.

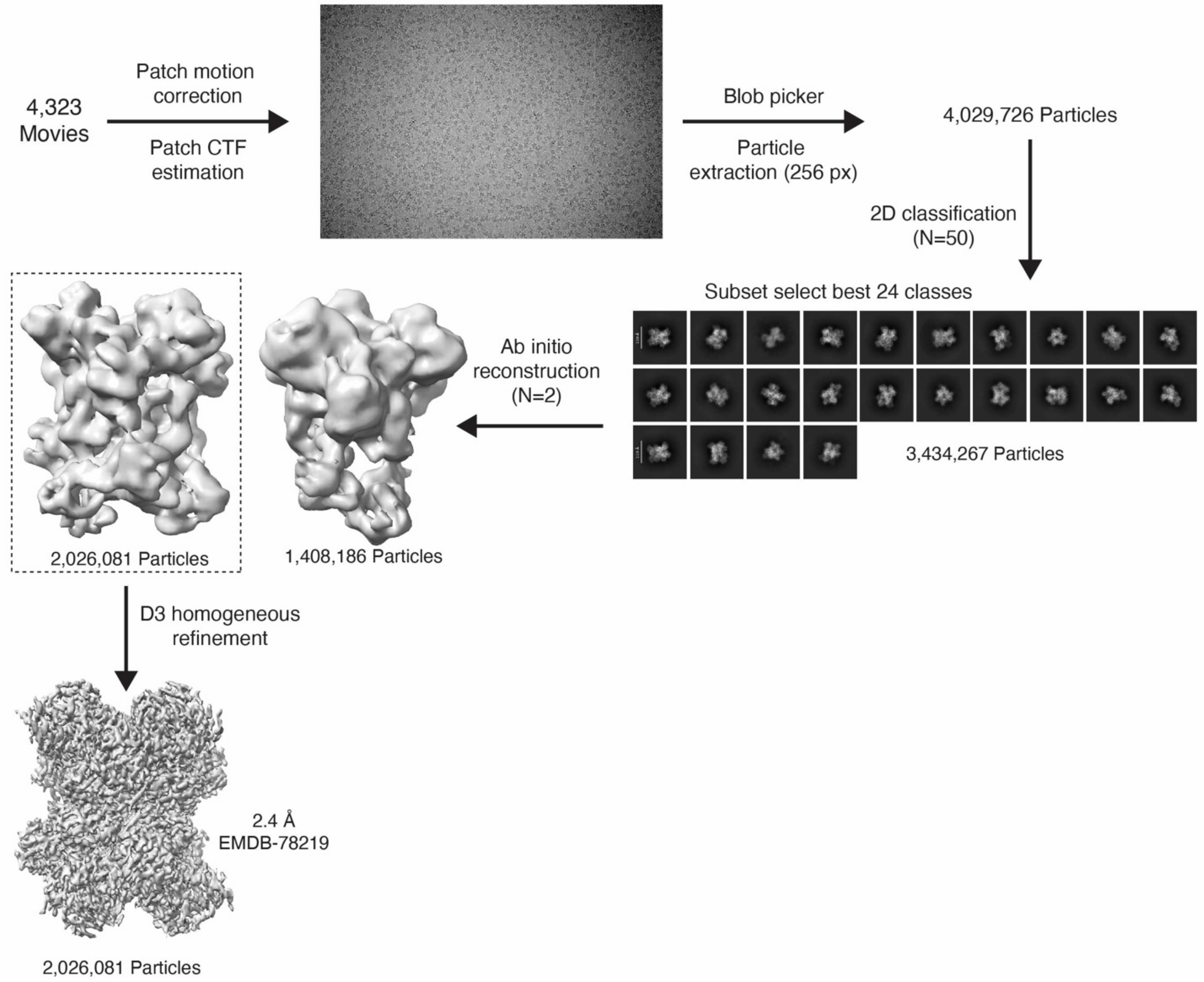


**Supplementary Figure 3.** Cryo-EM data processing in CryoSPARC (6) of the apo-IBV nsp15 dataset collected on the Talos Arctica (ThermoFisher), starting with 4,323 movies. From 4,029,726 particles extracted, 3,434,267 particles were classified in 2D class averages and used for Ab initio reconstruction. From the two classes produced during Ab initio reconstruction, one final map was refined to a resolution of 2.4 Å.

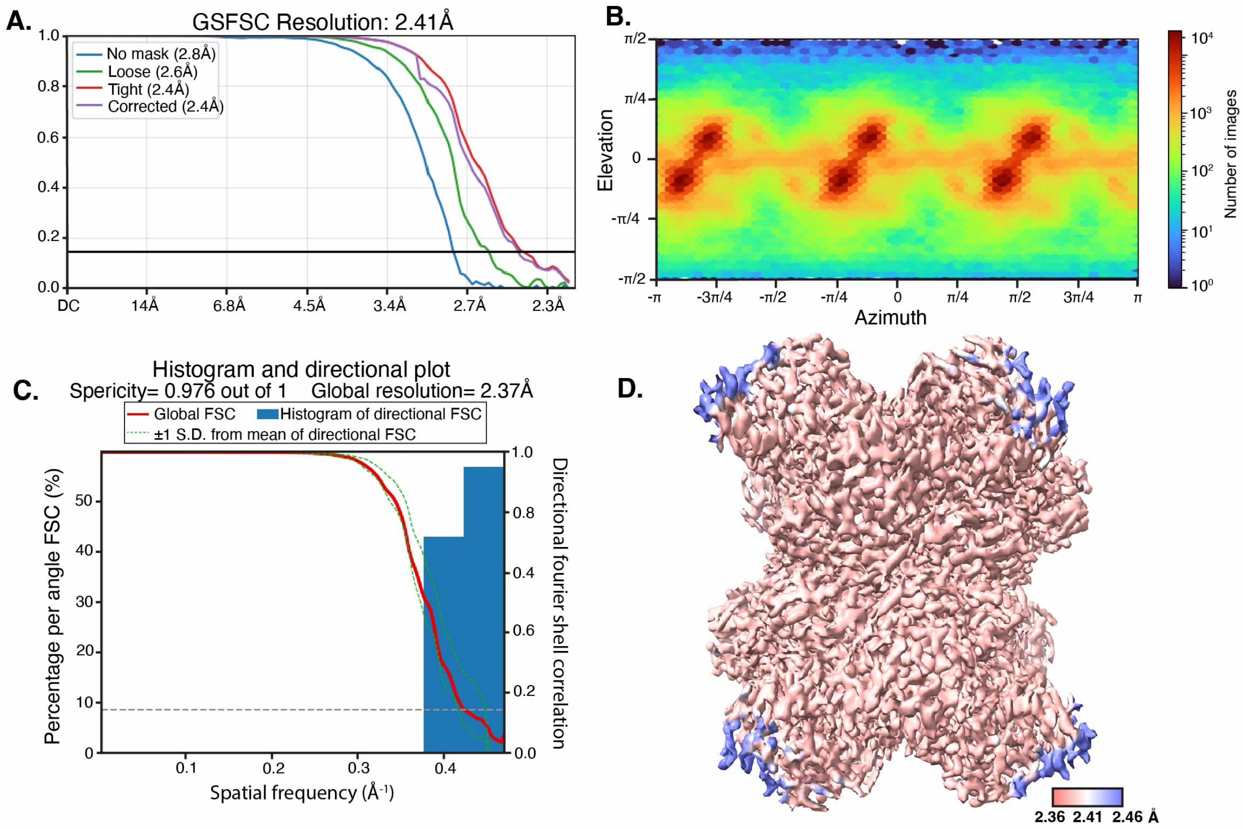


**Supplementary Figure 4.** Apo-IBV nsp15 map validation. (**A**) The GSFSC output from CryoSPARC shows a final map resolution of 2.4 Å. (**B**) Angular sampling of the projections was determined using CryoSPARC. (**C**) The 3DFSC output shows a sphericity of 0.98. (**D**) The local resolution map shows a majority of the structure is resolved to a narrow resolution range.

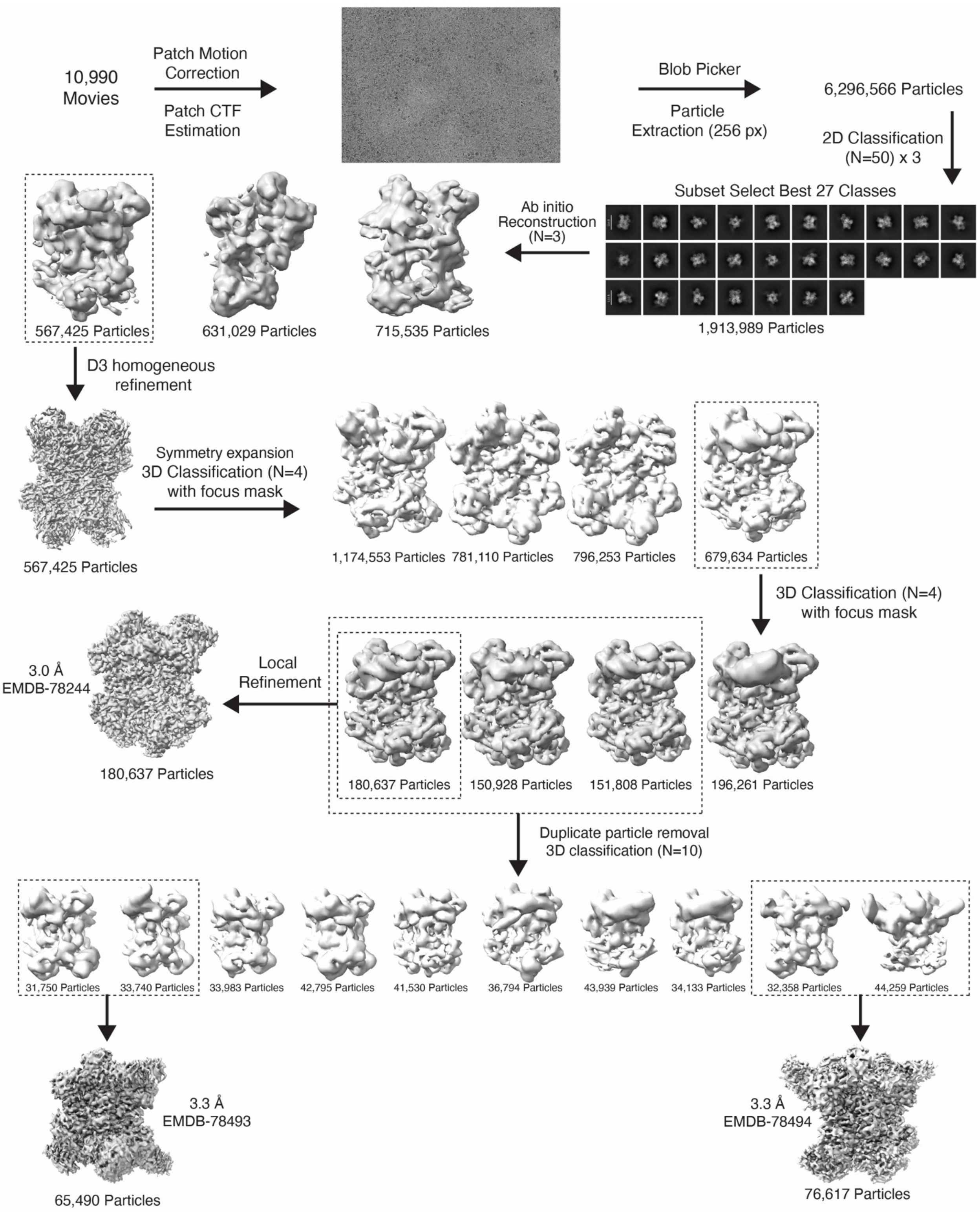


**Supplementary Figure 5.** Cryo-EM data processing in CryoSPARC (6) of the dsRNA bound-IBV nsp15 dataset collected on the Talos Arctica (ThermoFisher), starting with 10,990 movies. From 6,296,566 particles extracted, 1,913,989 particles were classified in 2D class averages and used for ab initio reconstruction. From the three classes produced during ab initio reconstruction, one class was homogeneously refined and the corresponding particle set was symmetry expanded. A mask including one protomer and bound dsRNA was used in combination with the symmetry expanded particles to resolve the dsRNA density. After 3D classification and local refinement, a final map containing one dsRNA bound was obtained at 3.0 Å. From the 3D classified volumes, two maps which contained two dsRNAs bound to nsp15 were resolved to 3.3 Å.

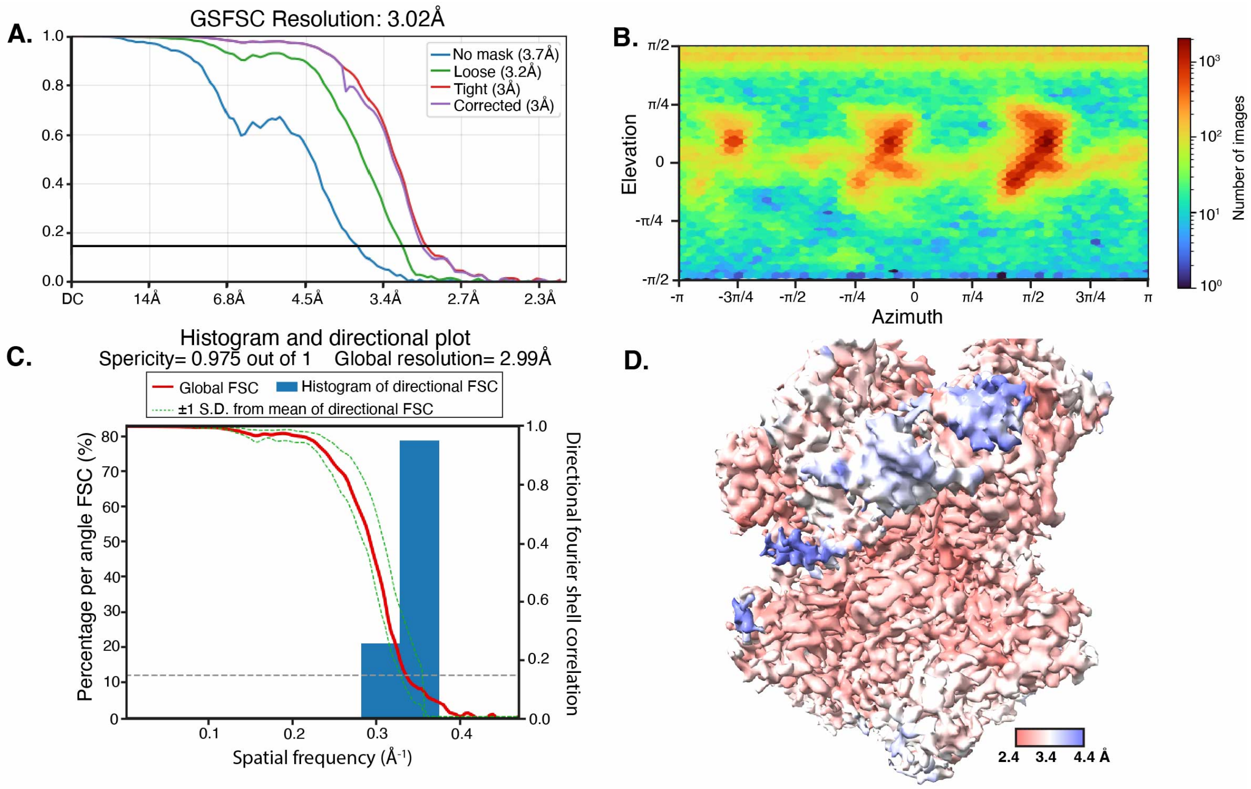


**Supplementary Figure 6.** Single dsRNA bound-IBV nsp15 C1 reconstruction validation. (**A**) The GSFSC output from CryoSPARC indicates a final resolution of 3.0 Å. (**B**) The angular sampling of the projections shows a distribution of views contributing to the final reconstruction. (**C**) The 3DFSC indicates a sphericity of 0.98. (**D**) The local resolution map shows that the dsRNA has lower resolution compared to the rest of the structure.

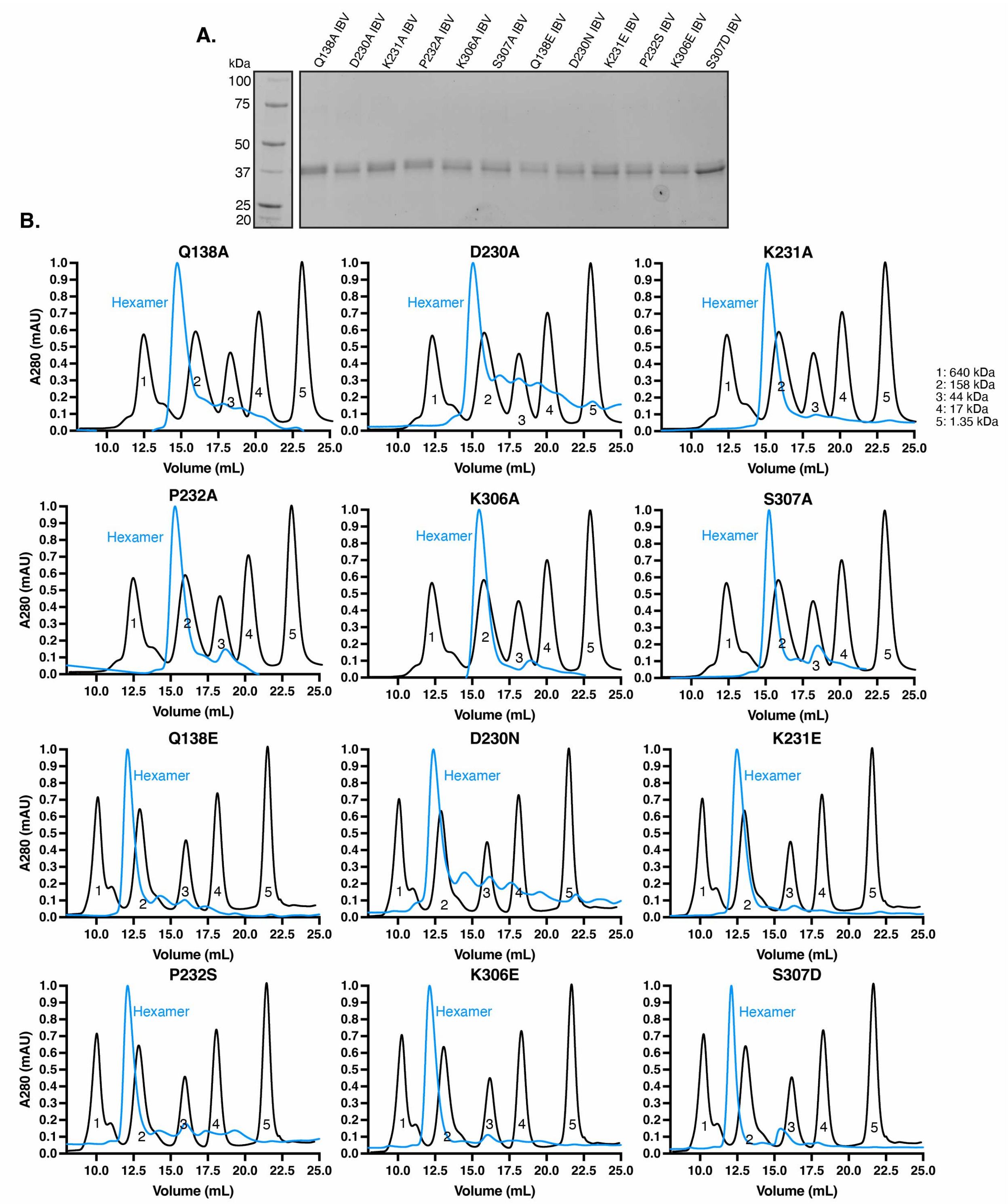


**Supplementary Figure 7.** Purification of nsp15 proteins. (**A**) The SDS-PAGE of mutant IBV nsp15 proteins, post-size exclusion chromatography shows a pure protein at 38 kDa. (**B**) The size exclusion chromatography profiles of the IBV mutant proteins shows the main peak as hexameric protein.

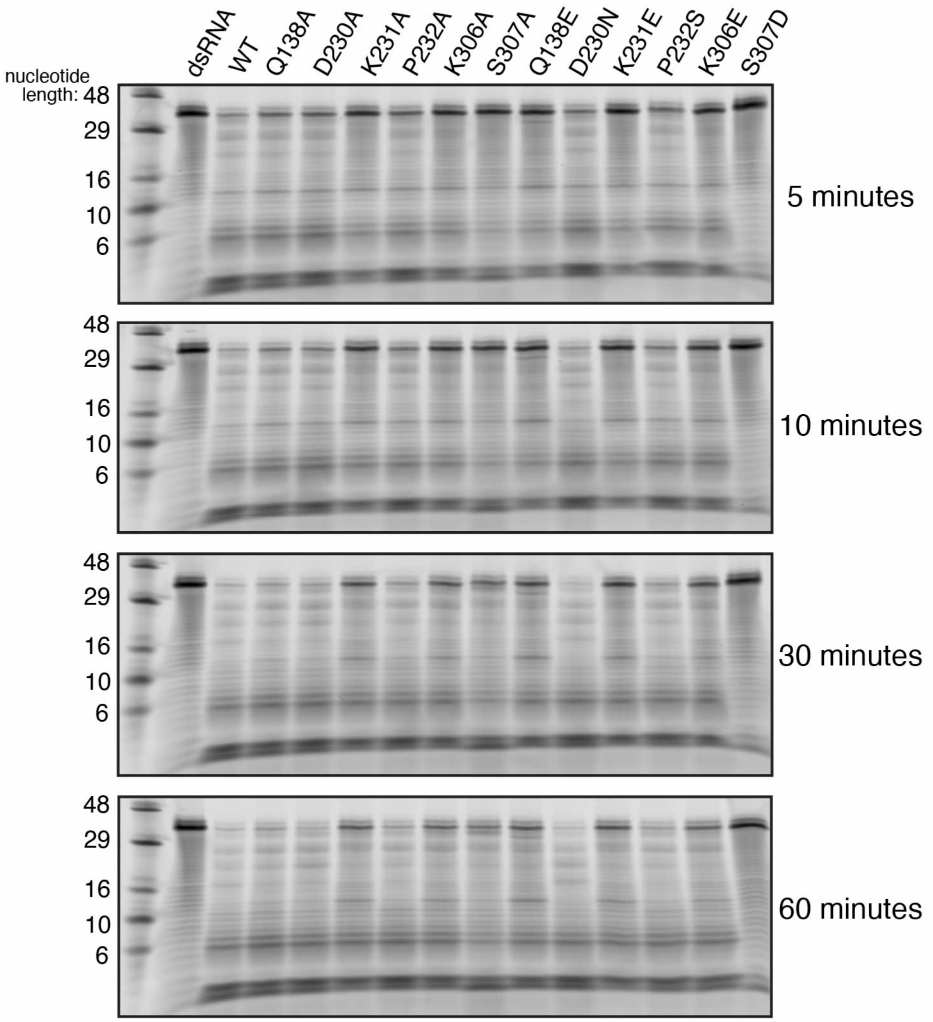


**Supplementary Figure 8.** Gel analysis of dsRNA cleavage by IBV WT and mutant nsp15s at four time points, shows varying dsRNA degradation by nsp15.

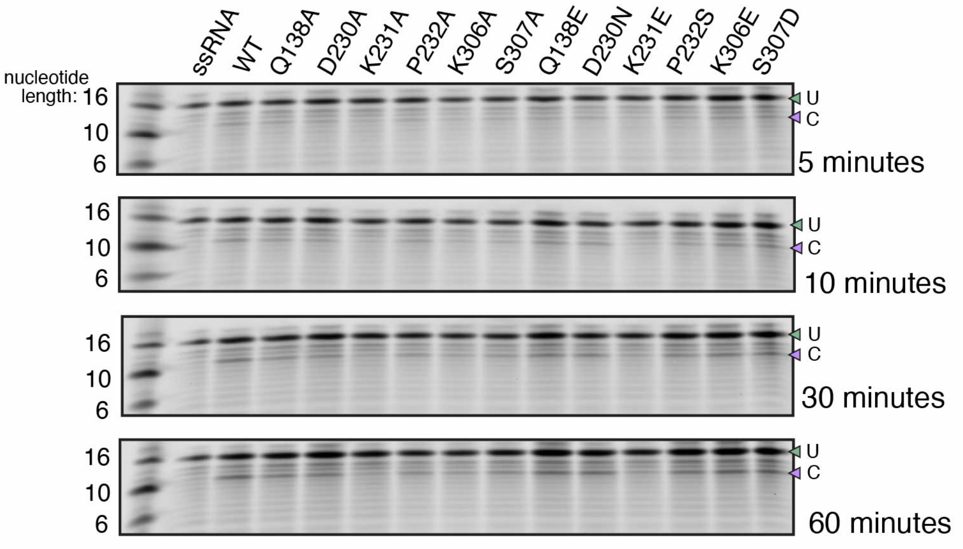


**Supplementary Figure 9.** Gel analysis of ssRNA cleavage by IBV WT and mutant nsp15s at four time points shows a single cleavage product of 13 nt (U: uncleaved band and C: cleaved band).

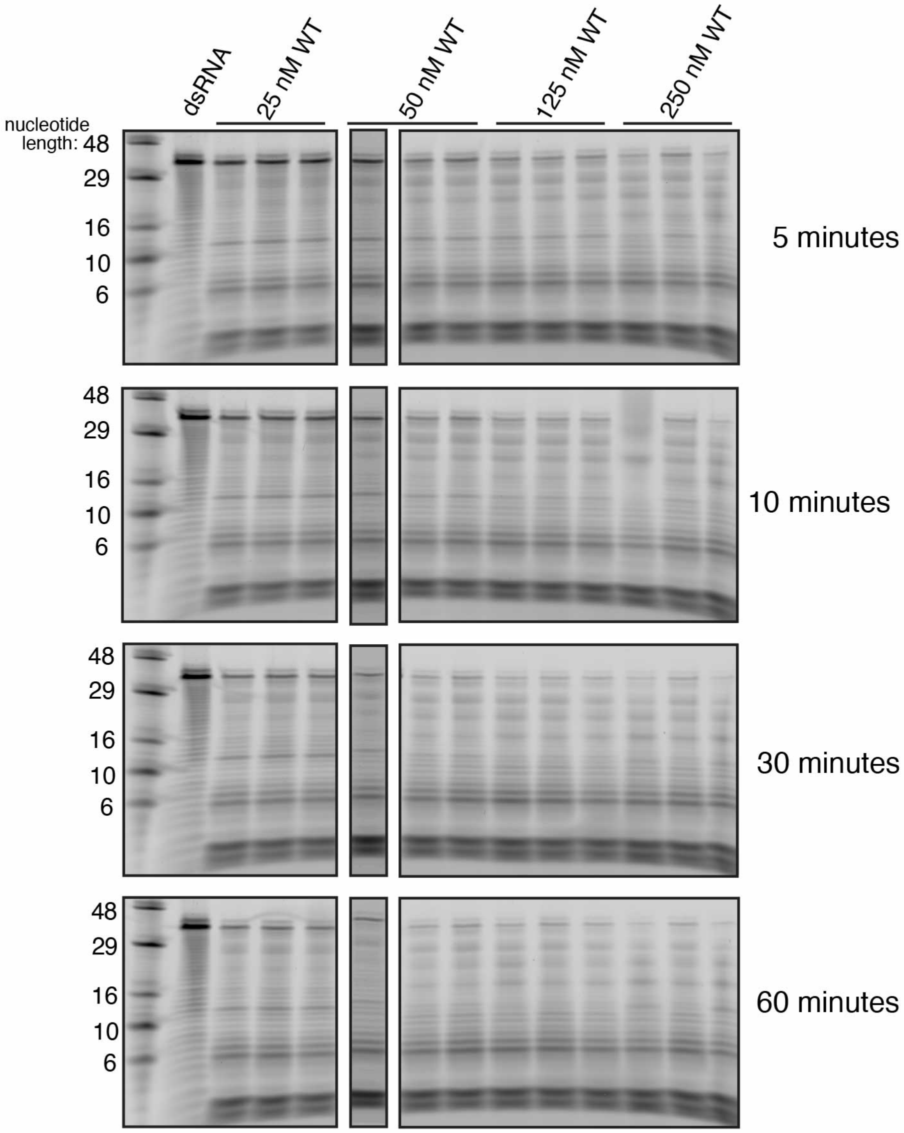


**Supplementary Figure 10.** Gel analysis of dsRNA cleavage with varying amounts IBV WT nsp15 at four time points shows multiple cleavage products.

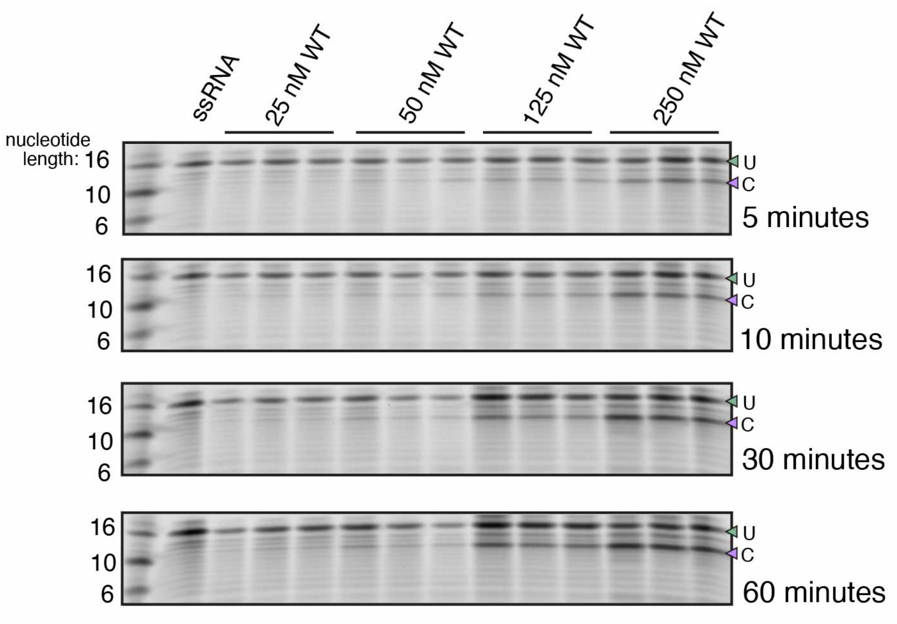


**Supplementary Figure 11.** Gel analysis of ssRNA cleavage with varying amounts IBV WT nsp15 at four time points displays a single cleavage product at 13 nt U: uncleaved band and C: cleaved band).

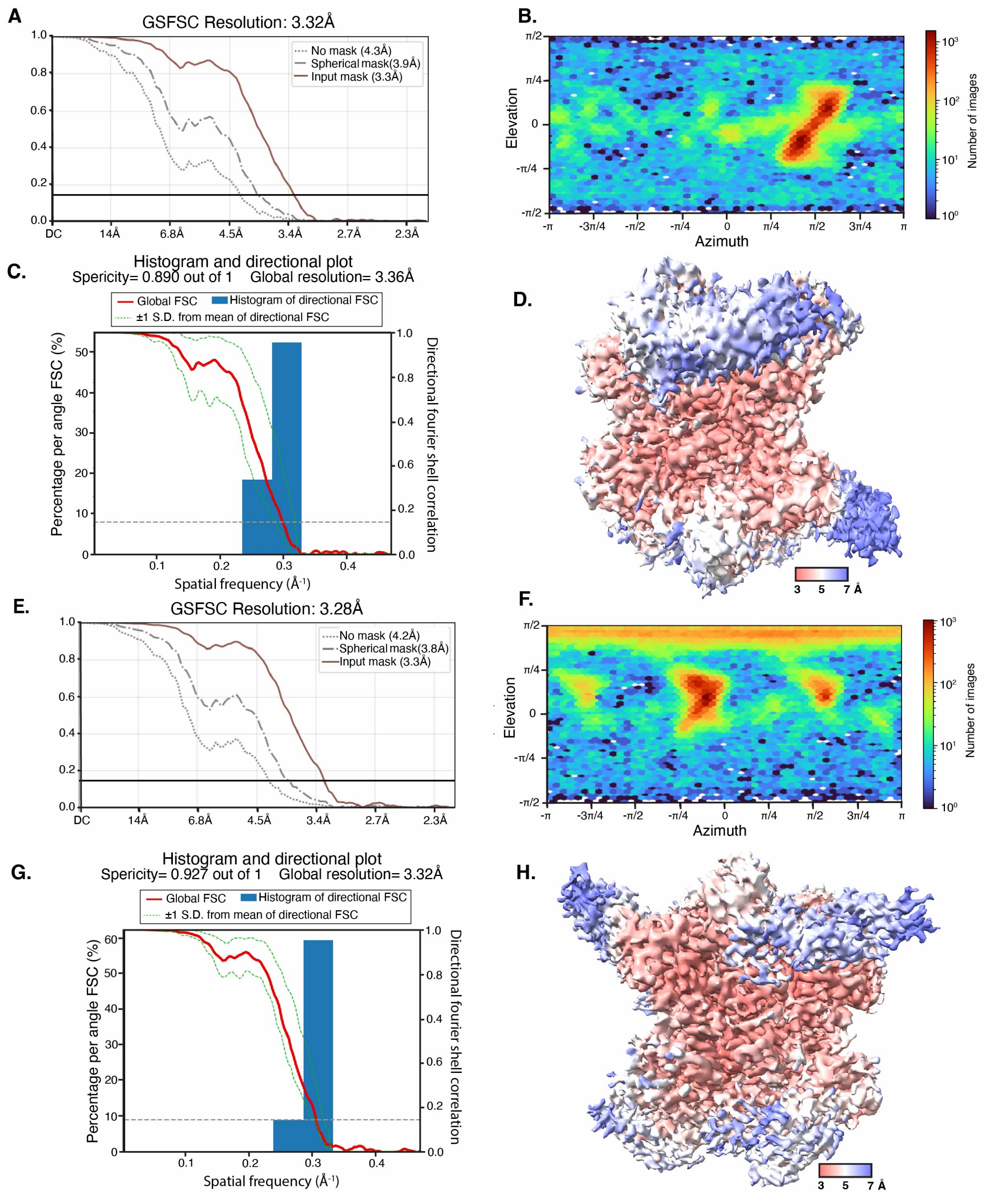


**Supplementary Figure 12.** Dual dsRNA bound-IBV nsp15 map validations. (**A**) The GSFSC output from CryoSPARC shows a final resolution of 3.3 Å for the map with a dsRNA bound to separate trimers. (**B**) The angular sampling of the projections displays a majority of particles in a predominant orientation. (**C**) The 3DFSC output shows a sphericity of 0.89. (**D**) The local resolution map shows higher resolution for the protein compared to the dsRNA. (**E**) The GSFSC output from CryoSPARC has a final resolution of 3.3 Å for the map with dsRNAs bound to the same trimer. (**F**) The angular sampling of the projections shows a small collection of predominant views approximately 120˚ apart. (**G**) The 3DFSC shows a sphericity of 0.93. (**H**) The local resolution map is similar to panel (D), where better resolution is seen for the protein compared to the dsRNA density.
